# Perception of Mangrove Ecosystem Conservation: A Case of Silvofishery in East Kalimantan, Indonesia

**DOI:** 10.1101/2025.03.25.645347

**Authors:** Aji W. Anggoro, Yustina Octifanny, Dini Aprilia Norvyani, Nabilla Dina Adharina, Muhammad Ilman, Topik Hidayat, Mariski Nirwan, Basir, Vabian Adriano, Rahmadi Muis, Muhammad Danie Al Malik, Andi Trisnawati

## Abstract

This research aims to explore farmers’ perceptions of mangroves, the impact of mangroves availability on shrimp aquaculture production, and the relationship between mangroves and aquaculture production among traditional shrimp aquaculture farmers in Berau. This study utilized a mixed-method approach to investigate farmers’ perceptions of mangroves and their contribution to aquaculture production. Quantitative data was collected through surveys targeting traditional shrimp farmers across six villages in Berau during September-October 2023, covering 619 ponds (64.01% of the total 967 ponds). The data analysis focused on mangrove availability, positioning within ponds, and intensity surrounding them, using two-way MANOVA to explore the influence of mangroves on shrimp production, preceded by t-tests and one-way ANOVA to compare production across different pond characteristics. The quantitative analysis was enriched by qualitative data, obtained through 34 semi-structured interviews utilizing purposive sampling, aimed to capture diverse perspectives on the mangrove ecosystem’s role in aquaculture. Our findings suggest that the integration of mangrove and shrimp aquaculture holds potential, but must address shrimp farmers’ concerns regarding livelihood continuity and technical challenges such as ensuring suitable mangrove species and managing stable pond conditions. By exploring the community’s views on the degradation of integrated mangrove and shrimp aquaculture and its impacts, this study develops strategies to engage shrimp farmers in mangrove restoration efforts while supporting their livelihoods.

## Introduction

Between 2001 and 2020, Indonesia lost 193,367 hectares of mangroves driven by deforestation (148,727 ha), commodities (38,669 ha), erosion (3,660 ha), and other factors (2,306 ha) [1]. Key factors that drive mangrove loss in Indonesia are the conversion of mangrove forests into aquaculture, oil palm, and other land uses [2]. According to Murdiyarso et al.[3], the mangrove forest that has been converted into aquaculture has the least total average ecosystem carbon stock (579 Mg C ha−1), in comparison to degraded mangrove (717 Mg C ha−1), regenerated mangrove (890 Mg C ha−1), and undisturbed mangrove (1061 Mg C ha−1).

Between 2012 and 2030, Indonesia expects a 7% growth in aquaculture production on approximately 26 million hectares of land, mainly on existing wetland and mangrove areas for shrimp production [4]. The impact of mangrove conversion to shrimp aquaculture will harm the climate mitigation function of mangrove forests, such as hosting coastal species, coastal protection, deposing sediments, and storing pollutants [5]. The loss of mangroves to shrimp aquaculture land uses also emits more carbon to the atmosphere. In addition, the conversion of mangroves to shrimp ponds results in the loss of 226 years of soil carbon accumulation in natural mangroves, and each kilogram of Tiger Shrimp is equivalent to almost 1,000 liters of gasoline consumed by automobiles [6].

Indonesia has proposed an ambitious rehabilitation target of 600,000 ha of mangrove by 2024 in nine provinces–North Sumatra, Riau, Riau Islands, Bangka Belitung, West Kalimantan, East Kalimantan, North Kalimantan, Papua, and West Papua–according to [1]. The conservation effort has been complicated by the land regulatory misfit among the governing institutions that manage mangroves nationally in Indonesia [7]. The mangrove forest is designated into three land statuses: protected forest area, production forest area, and other uses (APL or *Area Penggunaan Lainnya*); it comprises state and private lands that make government, private actors, and community manage the mangroves and activities above them [8]. Most of the mangrove losses are emerging on three different land uses with different extents: 91,545 ha in APL, 53,131 ha in production forest, and 48,691 ha in protected forest [1]. Until the 2020s, more than 20% of mangroves in APL land are in critical condition, making it important for the government actors to collaborate with the local community and private landowners to conduct mangrove conservation and sustainable management while preventing livelihood disturbance to landowners and coastal communities [8].

East Kalimantan Province can potentially restore 20% of the national restoration target encompassing 40,000 ha area in six districts, including Banyuasin, Bulungan, Tana Tidung, Paser, Berau, and Nunukan [1]. Similar to the challenge on the national level, mangrove restoration in East Kalimantan’s Districts is difficult to implement owing to the community’s and landowners’ shrimp aquaculture development. Following the Southeast Asia Shrimp Boom since the 1980s, the mangrove in East Kalimantan, particularly in the Berau District, is mainly under the custody of people whose main livelihood is the traditional fish and shrimp aquaculture [9].

The aquaculture community in Berau has passed down knowledge and experiences in aquaculture management, but they have a low awareness level and participation rate in community-based mangrove management [10–11]. According to Lukman et al. [10], the aquaculture farmers in Berau have a limited understanding of mangrove benefits, and their experience with aquaculture activities narrowly defines their perception of mangroves only as erosion prevention and local fish and shrimp species nursery. On the other hand, the friction between the mangroves has been cast in a negative light due to their ecosystem disservices to the shrimp production, such as polluting the ponds, contributing faster pond sedimentation, irregulating water temperature, and being a place to host predators without insufficient drainage and management system [5]. Regardless of the mangrove-aquaculture contention in the Berau District, it poses an opportunity for small-scale mangrove restoration. Small-scale restoration of low-productivity or abandoned aquaculture ponds due to disease, sub-optimal water management, and exposed coastline to erosion and extreme tides hold the largest restoration opportunities if accumulated, according to Lovelock et al [12].

In Berau District and other areas experiencing the mangrove-aquaculture tension, mangrove forest conservation and restoration management depend on the farmers’ perception of mangroves, both service and disservice, in conjunction with their aquaculture production. Using case study research in Berau District, East Kalimantan, Indonesia, this study provides an evidence-based restoration opportunity by understanding the perception and livelihood-driven barriers in traditional and small-scale shrimp aquaculture communities to partake in small-scale mangrove conservation. We want to explore three questions concerning traditional shrimp aquaculture farmers in Berau: What are farmers’ perceptions of mangroves and their future aspirations towards the remaining mangroves? What is the impact of the difference of mangroves availability on the production of shrimp aquaculture? How does mangrove affect aquaculture production?

By understanding traditional shrimp aquaculture-induced mangrove degradation from a community perspective and their livelihood, this study hopes to map a meaningful strategy to engage the shrimp farmer community in mangrove restoration without disturbing their opportunity for a dignified livelihood. Combined with addressing socioeconomic inequality, negotiating land tenure insecurity, realigning policy regulation, maintaining farmers income while restoring the mangrove, providing alternative livelihood, and building community-led mangrove restoration management [13], this study hopes to move the small-scale restoration agenda beyond mangrove planting, conserving remaining mangrove stands, and restoring critical mangrove areas.

## Materials and methods

### Location

Berau District is administratively located in East Kalimantan, Indonesia, with its territory consisting of inland, urban, and coastal areas. Berau District has a 36.962,37 km² total area with 39.85% of it being water area, or about 14,729.86 km² [22]. Although Berau is not a small island district per se, the area is considered a small and remote island. Berau District is part of the Coastal Conservation Areas and Small Islands Derawan Islands and Surrounding Waters (*Kawasan Konservasi Pesisir dan Pulau-Pulau Kecil Kepulauan Derawan dan Perairan Sekitarnya* - KKP3K-KDPS), with mangrove forests covering an area of around 80,000 hectares, the largest in East Kalimantan [23]. KKP3K-KDPS has a 22,213 ha mangrove ecosystem or about 25% of the mangrove area in Berau District. KKP3K-KDPS is a part of coastal and marine conservation areas targeted to be conserved by the government in 2030 where the mangrove forests are in a natural state for the most part [24]. Berau’s mangrove ecosystem is considered as good in that it provides nutrition for fish breeding as well as tourism potential [11].

Our study area in this research involved six villages spread across three sub-districts on the coast of Berau District (Fig 1). These six villages are dominant in terms of mangrove presences and shrimp aquaculture. These villages are Pegat Batumbuk Village in Pulau Derawan Sub-District; Suaran Village, Sukan Tengah Village, and Pilanjau Village in Sambaliung Sub-District; and Tabalar Muara Village in Tabalar Sub-District.

**Fig 1.**
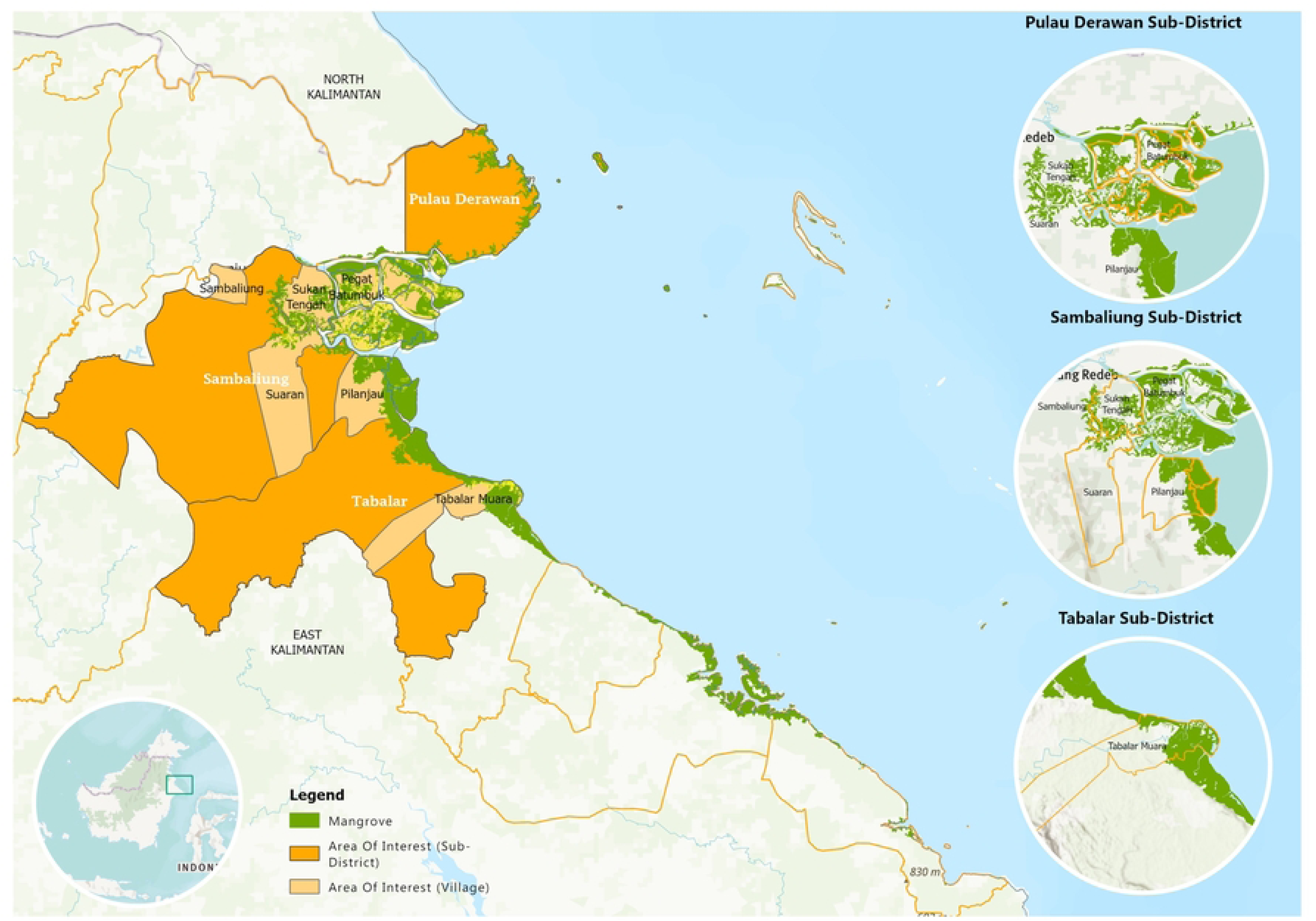
The map of Berau District and surveyed villages within three sub-districts

### Study design

Prior studies on perceptions towards the environment suggest that communities’ attitudes, views, and perceptions play a pivotal role in shaping environmental management and ecosystem-based livelihood strategies, including the success of conservation efforts [15–16]. A study by Firdaus et al. [17] revealed that different perceptions between the fishery community and shrimp farmers determine their attitudes towards improving mangrove conditions. Conversely, people’s livelihood and demand form their viewpoint on ecosystem services.

This study employed a mixed-method approach with a concurrent triangulation design to understand the farmers’ perception of mangroves and how mangrove contributes to aquaculture productivity. A mixed-method approach allows the researcher to gain deeper meaning and comprehensively understand the phenomenon studied [25–26]. Morse [26] claimed that in mixed-method research, the study must have one core component, either quantitative or qualitative.

### Quantitative method

The quantitative data for this study was collected through surveys targeting traditional shrimp farmers in September-October 2023. Preliminary mapping identified 967 ponds across six villages, but only 619 active ponds (64%) were surveyed, including locations such as Gurimbang (5 ponds), Pegat Batumbuk (518 ponds), Suaran (44 ponds), Sukan Tengah (7 ponds), Tabalar Muara (44 ponds), and Pilanjau (1 ponds). The detailed data related to total ponds for each village that had been collected is in S1 Table.

Before the pond survey, we trained ten enumerators for four days. They were trained to conduct surveys using ODK Collect, in-depth interviews, data management, research ethics, and safety and security. The quantitative data collection uses an ODK collection application installed into the enumerators’ smartphones. After the training, the enumerators conducted pilot testing. There was a feedback mechanism in which the questionnaire was adjusted based on the pilot test experiences. The adjustment reduced the number of questions and replaced complex terminologies with local vocabulary. There were weekly quality control activities between the researchers and the enumerators.

Farmers provided responses based on each individual pond they owned or managed, leading to multiple contributions from single farmers. Male farmers dominated in this region [27] with represented 94.83% (587 ponds) and female farmers 15.17% (32 ponds). Most farmers were aged 40-49 and had primary school education. The average pond area was 16.44 hectares, with 91% (568 ponds) cultivating shrimp and other commodities like milkfish and crab.

Due to the proximity of mangroves to aquaculture operations in Berau Regency, 333 farmers incorporate mangroves within their ponds, while 286 farmers’ ponds lack mangroves within their confines. S2 Table displays the detailed profile of respondents involved in a questionnaire survey. Among these, 83% (516 ponds) included mangroves. The questionnaire covered farmers’ profiles, pond and livelihood details, aquaculture production and inputs, and perceptions towards mangroves. The research also collected data on farmers’ perceptions of mangrove benefits and associated issues, the reasons why farmers are maintaining mangrove stands, and their future aspirations regarding mangroves.

### Qualitative method

The qualitative method explores the diversity of knowledge that can enrich, complement, or contradict the quantitative data. An in-depth interview is the primary qualitative data collection method. It is designed to explore the following information: 1) Respondents’ backgrounds and credentials, 2) Pond History, 3) The state of farmers’ aquaculture productivity, 4) Farmers’ testimony on the reasons for an increase or a decrease in productivity, 5) Farmers’ consideration for conserving natural mangroves, exploiting natural mangroves, or adopting integrated mangrove-aquaculture system, and 6) Farmers’ perspective on the connection between mangrove conservation and their aquaculture production. For sampling or participant selection, the authors employed a purposive sampling strategy called maximum variation sampling to capture diverse experiences and points of view and select respondents along various characteristics that may have relationships with their perceptions of and experiences with the mangrove ecosystem. The 34 semi-structured interviews were performed with the participant’s key characteristics:

● Integrated Mangrove-Shrimp Aquaculture (IMSA). From the preliminary survey, at least four farmers have fully adopted the IMSA.
● Representation of villages. The interviewed representation of respondents from each village. Owing to unequal shrimp pond area, number, and ownership in and across each village, the sample of informants is weighed based on the distribution of the pond and ownership.
● Diversity in the background of farmers and farm workers. We interviewed shrimp middlemen and/or collectors, while the rest of the respondents are traditional shrimp farmers. The farmers vary based on gender, pond ownership (pond owners and workers), and income level (low and high).

The interview transcripts were analyzed using coding clustered into three clusters: shrimp farmers-pond background, middleman background, and mangrove experience/perception (see S3 Table).

### Data analyses

The quantitative analysis method employed is two-way MANOVA (Multivariate Analysis of Variance). MANOVA expands upon ANOVA by investigating how one or more categorical independent variables influence multiple continuous dependent variables [28], and two-way MANOVA is a statistical technique that extends the principles of ANOVA to assess the effects of two categorical independent variables (factors) on two or more continuous dependent variables (variables). However, before performing MANOVA, this study first employed t-test and one-way ANOVA to describe and explore the production difference between different ponds’ characteristics in the context of mangrove presence. ANOVA is utilized to compare the production of ponds across various independent groups, each representing different characteristics. In this investigation, one-way ANOVA was conducted independently to examine and delineate the annual and cyclical production associated with each distinct independent variable listed in Table 1. Furthermore, prior to interpreting the comparative outcomes, Bartlett’s test for equal variance was applied. The shrimp production across nearly all groups exhibits unequal variance (See S4 Table), except total shrimp production in 2022 based on mangrove presence in external ponds (Chi2=0.558, p>0.05).

**Table 1.**
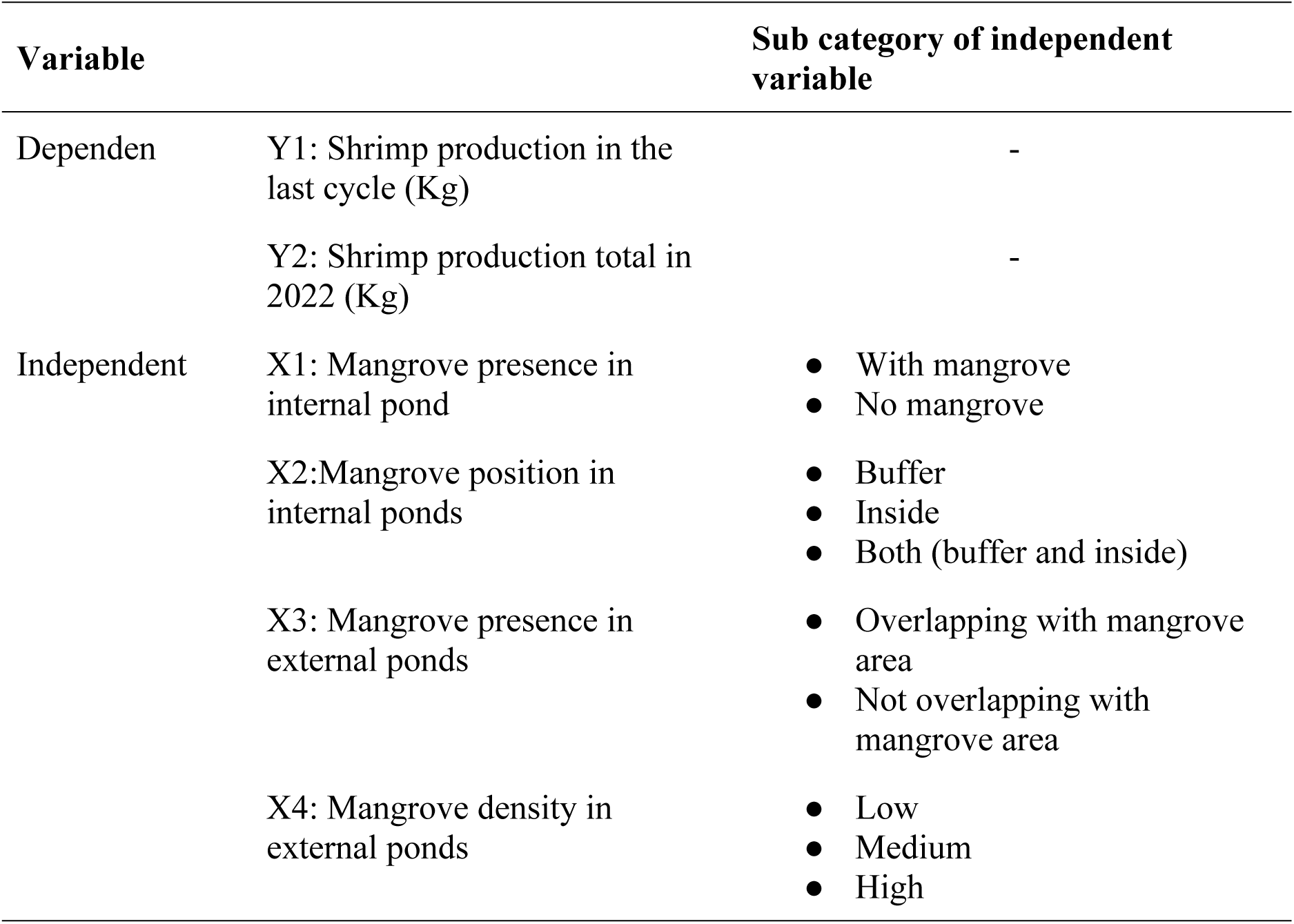
Dependent and independent variables.

| Variable |  | Sub category of independent variable |
| --- | --- | --- |
| Dependent | Y1: Shrimp production in the last cycle (Kg) | - |
|  | Y2: Shrimp production total in 2022 (Kg) | - |
| Independent | X1: Mangrove presence in internal pond | • With mangrove<br>• No mangrove |
|  | X2: Mangrove position in internal ponds | • Buffer<br>• Inside<br>• Both (buffer and inside) |
|  | X3: Mangrove presence in external ponds | • Overlapping with mangrove area<br>• Not overlapping with mangrove area |
|  | X4: Mangrove density in external ponds | • Low<br>• Medium<br>• High |

After performing ANOVA, two-way MANOVA was used to examine the interaction effect between the two independent variables, including mangrove presence in internal and external ponds, to the shrimp annual and cyclical production in kilogram units. The interaction effect tests whether the effect of one independent variable on the shrimp production variables depends on the levels of the other independent variables.

Two-way MANOVA was conducted twice. Firstly, it aimed to analyze the influence and interaction between the presence or absence of mangroves in internal ponds (X1) and the overlap or non-overlap with mangrove areas in external ponds (X3) on shrimp production in the last cycle (Y1) and total shrimp production in 2022 (Y2). This was conducted to all of the ponds’ samples from 619 ponds. Secondly, another two-way MANOVA was performed to examine the impact and interaction between the position of mangroves in internal ponds (X2) and the density of overlapping mangrove areas in external ponds (X4) on shrimp production in the last cycle (Y1) and total shrimp production in 2022 (Y2) in farmers’ ponds that have both mangroves in internal and external ponds. However, the second test was performed for only 164 ponds having both mangrove in internal and external location, hence the total ponds tested is smaller compared to the first one.

## Results

Although mangrove has been around people and farmers, the shrimp farmers have different perceptions of mangrove and its implication to the aquaculture production. From the survey, as seen in the Fig 3, 246 respondents believed that mangrove do not have any significant impact on pond aquaculture, 207 respondents agreed that mangrove leaf litter degrades the water quality, and 105 respondents believed that mangroves produce anti-nutrient content in the leaf litter that can decrease the production of shrimp (Fig 2).

**Fig 2.**
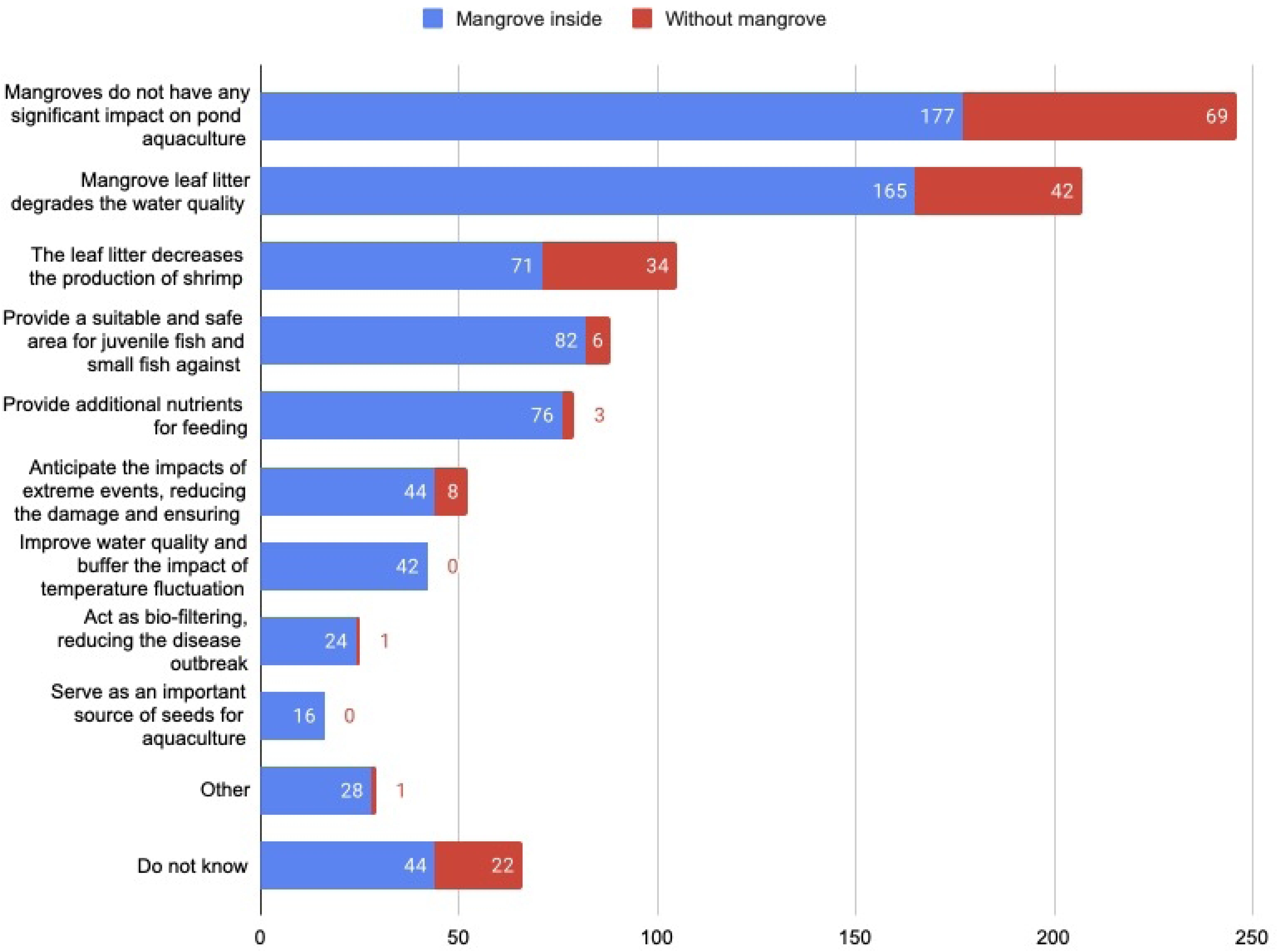
Pond-Level Data on Farmers’ Perception of Mangroves, Disaggregated by Mangrove Availability (Multiple Responses Allowed, n = 619 ponds)

**Fig 3.**
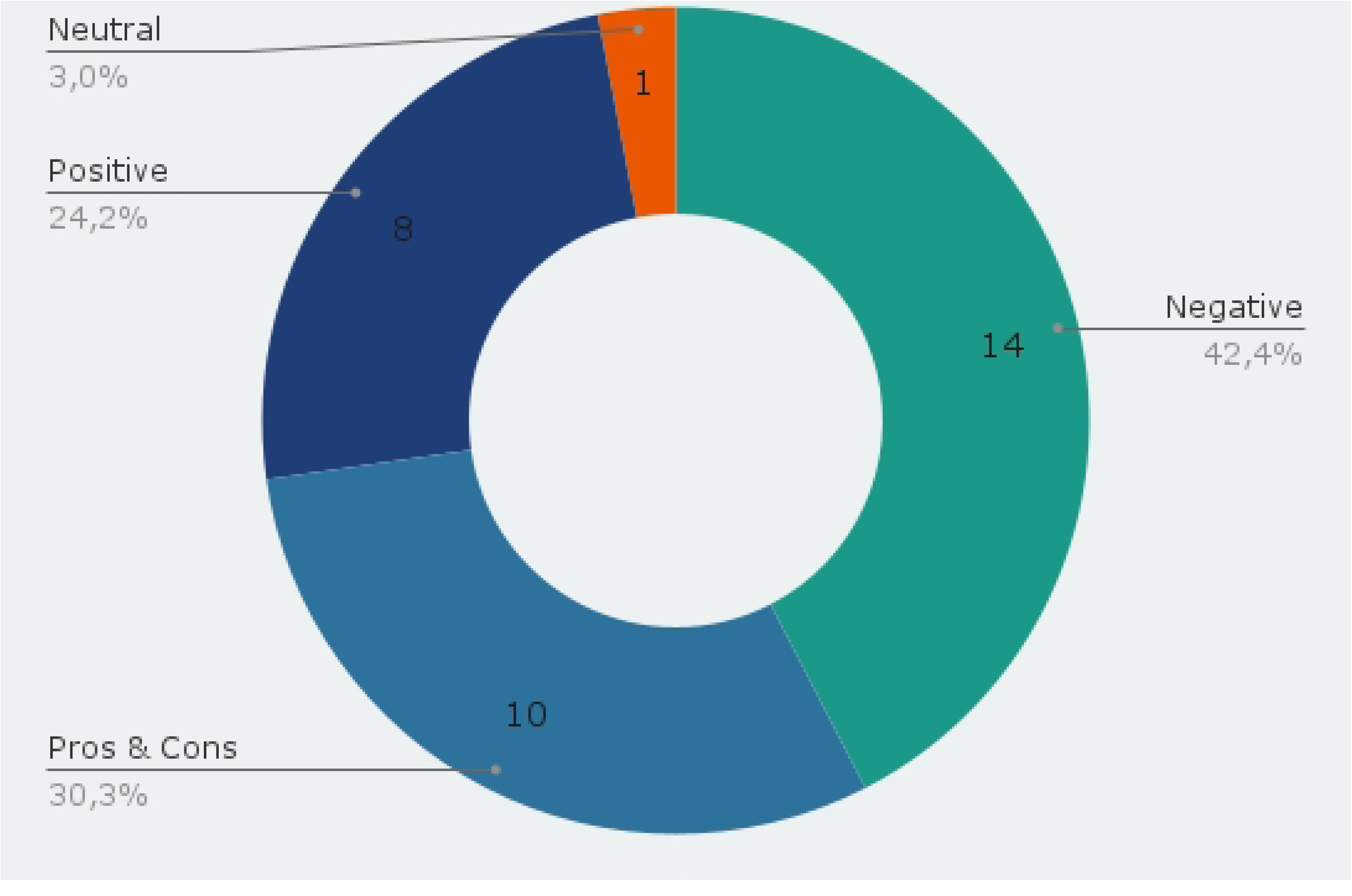
Farmers Perception to Mangrove from Interview (n = 33)

Similarly, from the 33 in-depth interviews, 24 respondents perceive that mangrove have a negative impact on the shrimp production by polluting the pond’s water with decomposition of leaf and branch litter (Fig 3). The water pollution also makes the shrimp prone to diseases. In addition, mangroves are also associated as the ground for shrimp predators to grow. For example, the community believe that shrimp larvae are being eaten by the small-size predators that live in mangrove roots or large shrimp are also being eaten by the beavers. Therefore, the majority of the farmers believe that having the pond that is clear from mangrove will boost their shrimp production. In addition, for the shrimp farmers, they can have more pond area if the pond is clear for mangrove. Mangrove also poses a practical challenge during the harvesting activities because it is not easy to harvest the shrimp in between mangrove roots.

From the interview, farmers believe that mangrove is a detrimental factor for declining shrimp production. However, from farmers’ testimonies, it is clear that there are other factors contributing to decreasing shrimp productivity beyond mangrove, including the quality of inputs (water, seeds, and feed), environmental conditions, diseases, and market conditions. Most of the interviewed respondents reported that their productivity is decreasing (59%), increasing (21%), consistent (9%), fluctuating (3%), while 6% do not know whether their productivity is increasing or decreasing.

With both positive and negative perception towards mangrove and its implication to reducing shrimp production, some farmers are still keeping mangrove inside or surrounding their pond. 120 ponds have mangroves inside it because the farmers think it is too hard and costly to clear the mangrove from the pond (Fig 4). There are 286 ponds with mangroves as buffers, where farmers claimed that mangroves play a role as a coastal defense, such as abrasion prevention, wave breaker, and flood protection. There are 51 ponds with mangrove inside them and 66 ponds with mangroves as buffers, where farmers believe that the mangroves have a role to protect their shrimp, juvenile fish, small fish, and crab from predators.

**Fig 4.**
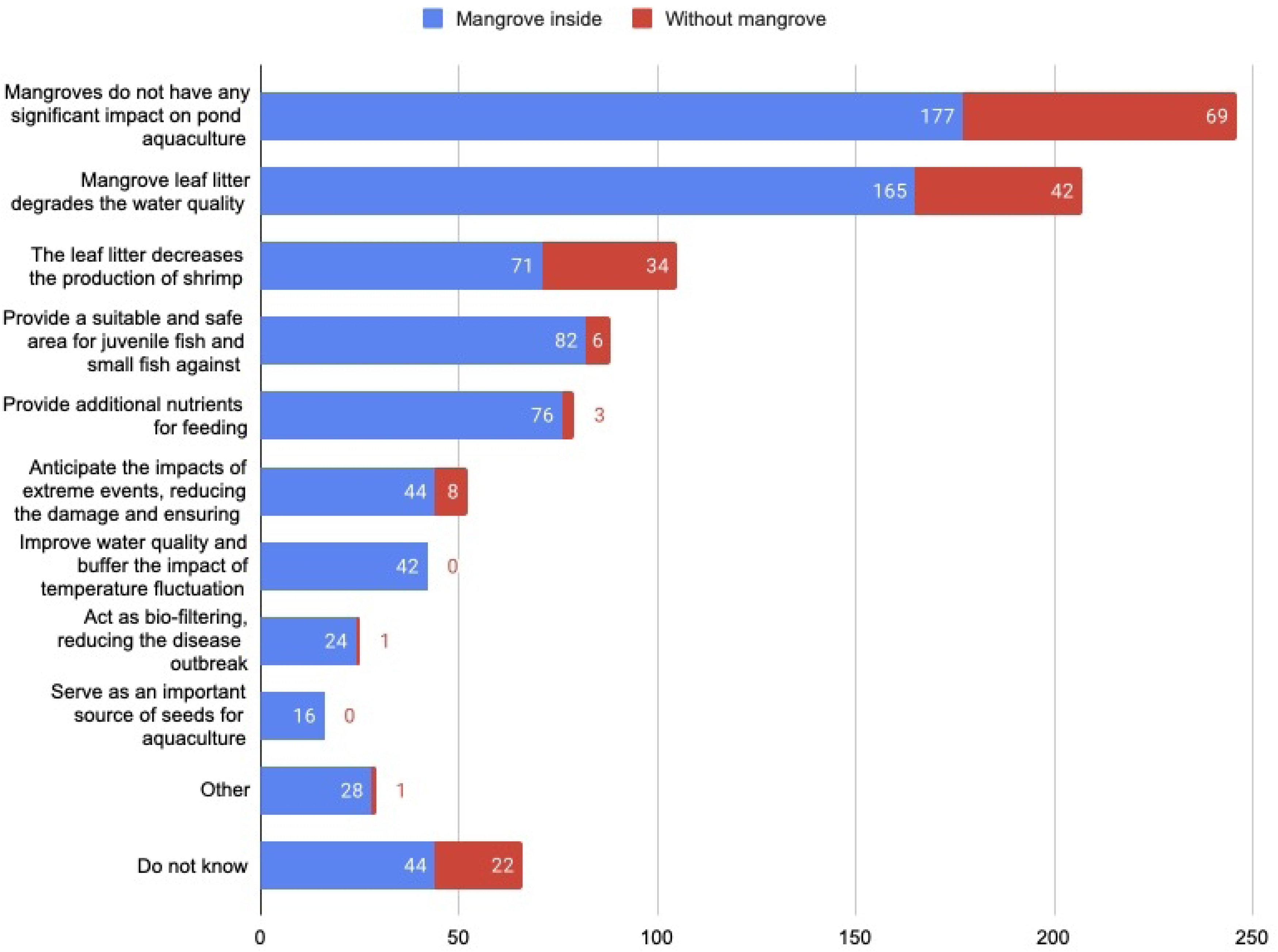
Pond-Level Data on Farmers’ Reasons for Keeping Mangroves Inside and Around Ponds (Multiple Responses Allowed, n = 516 ponds)

However, there is a disparity between farmers’ viewpoints regarding mangroves, reasons for decreasing productivity, and farmers’ aspiration for the future mangrove. It reveals a disparity in farmers’ viewpoints regarding the role of mangroves inside and around ponds (Fig 5). However, a number of farmers with mangroves within their ponds consider clearing or cutting them for aquaculture purposes. On the other hand, those utilizing mangroves as buffers perceive them as vital for coastal defense against erosion, large waves, and flooding. As farmers develop their perceptions of mangroves through personal interpretation, influenced by their surroundings and experiences in daily life and aquaculture, these perceptions will guide their subsequent actions concerning mangrove management.

**Fig 5.**
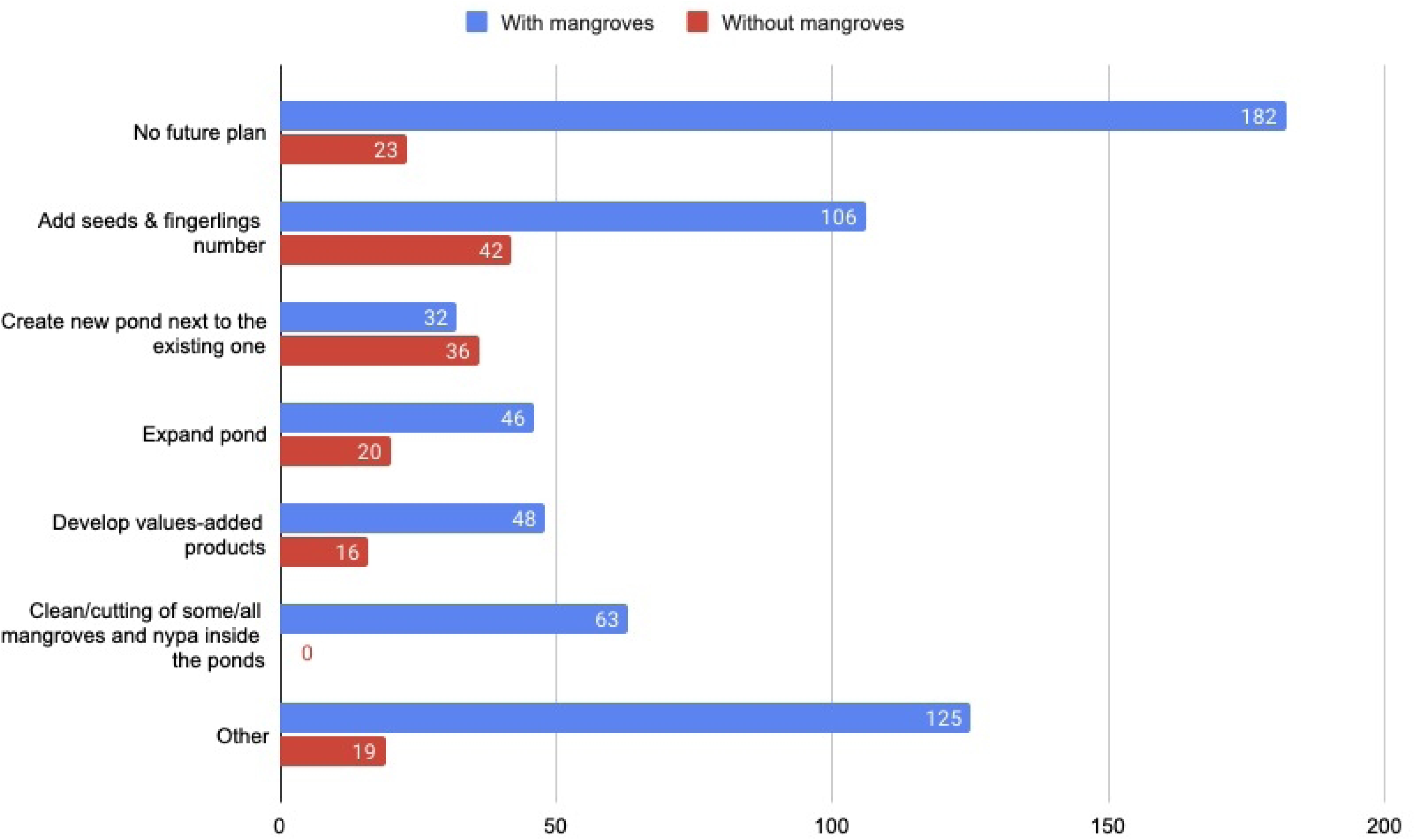
Pond-Level Data on Farmers’ Future Aspirations for Mangroves (Multiple Responses Allowed, n = 619 ponds)

Furthermore, the survey inquired about future plans concerning the mangroves. Out of 516 ponds with mangroves, 14 farmers intend to restore the existing mangroves in their ponds. Conversely, out of 516 ponds, 63 ponds were planned to have some or all of their mangroves and Nypa palms cleared or cut. Similar to the in-depth interview, although it is expensive to clear the land from mangrove, some farmers think it is necessary to cut the remaining mangrove for increasing shrimp production. Majority of the shrimp farmers intend to clear the mangrove in the middle of the pond, but they are keeping mangrove as a buffer of the pond so that their pond structure will be protected from land abrasion and extreme weather. Additionally, 31 ponds indicate other activities in the future, such as: divide their ponds, establish new ponds, sell their existing ponds, replant mangroves at reduced densities, enhance the mangrove structure, or transform the mangroves inside the ponds into palm oil plantations.

### 3.1. Mangrove presence on shrimp production (internal mangrove presence)

In Fig 6 and S5 Table, farmers having mangrove inside their ponds can produce 163 kg in the last cycle with a median 120 while farmers who do not have mangrove in their ponds can produce higher, 262.99 kg with a median 200 kg. Unequal t-test has proved that there is a significant difference between ponds with and without mangroves (t=-3.40, df= −114.469, dff= −99.38, p=0.000), with production from mangrove-free ponds exceeding those with mangroves by an average of 99.37 kg in the last cycle. In addition, farmers who incorporate mangroves into their ponds exhibit lower annual shrimp production with an average production of 526.05 kilograms and a median of 375 kilograms, whereas those without mangroves demonstrate higher production levels, averaging 783.20 kilograms, with a median of 500 kilograms. An unequal t-test indicates a significant difference in shrimp production (t = −2.62, df=118.97, diff=-257.14, p=0.009), with production from mangrove-free ponds exceeding those with mangroves by an average of 257.15 kg in 2022.

**Fig 6.**
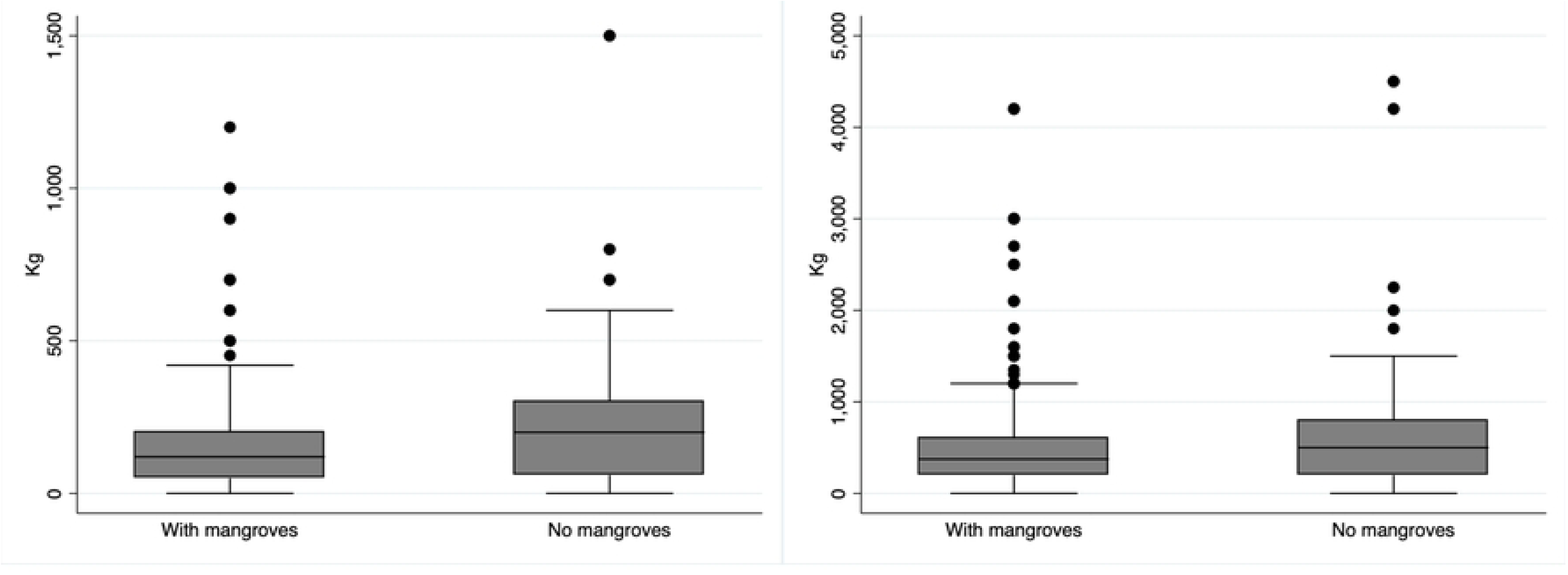
(Left) Total shrimp production in the last cycle based on internal mangrove presence, N=619 (Right) Total shrimp production in 2022 based on internal mangrove presence, N=619

Out of 516 ponds surveyed, 50 ponds integrated mangroves within the pond itself, 230 pounds utilized mangroves as buffers, and 236 ponds incorporate mangroves within the pond and as buffers. In Fig 7, distinct variations in the median of last cycle shrimp production and revenue are observed among ponds featuring mangroves within, as buffers, and in both capacities. Employing Robust ANOVA tests (see S6 Table), there is a significant difference shrimp production among the mangroves’ position in farmers’ ponds (Brown-Forsythe’s F(2, 76.57) = 3.67, p=0.029; Fisher’s F(2, 513) = 7.33, p=0.00; Welch’s F(2, 121.72)= 3.01, p=0.052).

**Fig 7.**
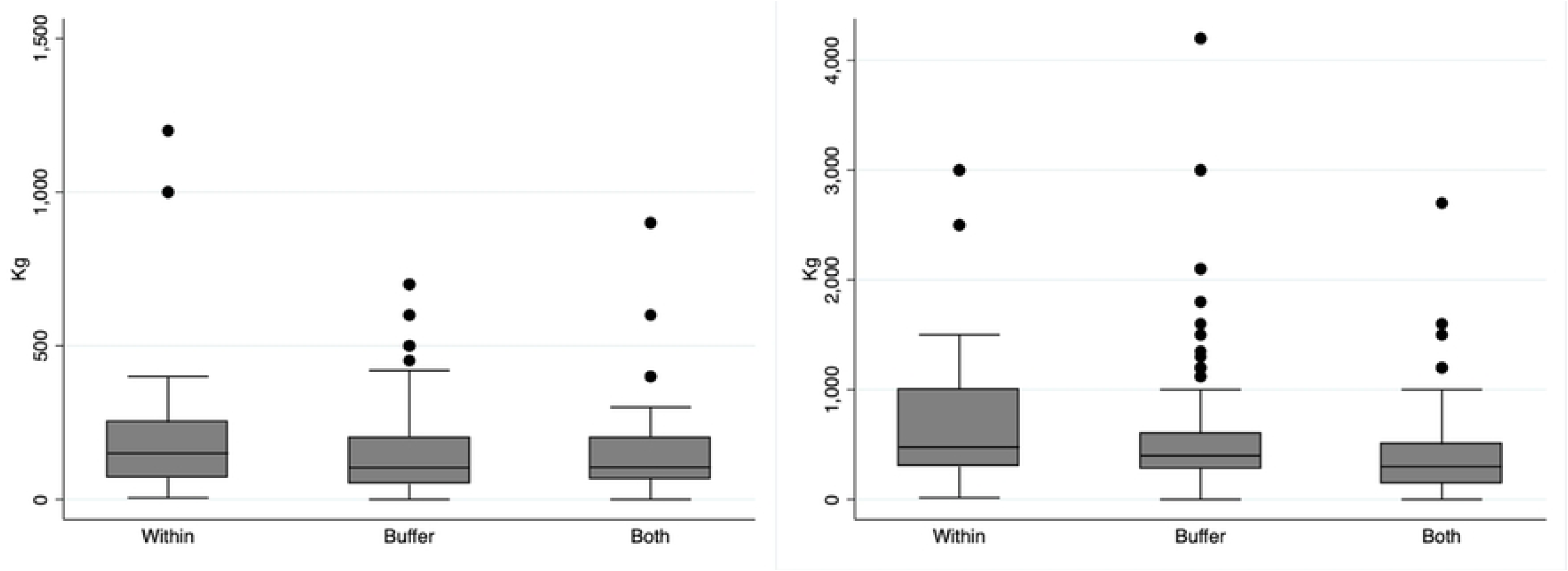
(Left) Total shrimp production in the last cycle based mangrove position, N=516 (Right) Total shrimp production in 2022 based on mangrove position, N=516

### 3.2. Mangrove presence on shrimp production (external mangrove presence)

Fig 8 depicts slight differences in production among ponds overlapped with mangrove area and not. The unequal t-test confirms that there is no significant difference among them (t=-0.47, df=617.81, diff=-7.89, p=0.632). The difference in cyclical production between ponds overlapping and not overlapping with mangrove areas is merely 0.47 kilograms. Similar with the cyclical production, subsequent t-tests reveal non-significant differences between shrimp production of ponds overlapping and not overlapping with mangroves area (t=-1.63, df=617, diff=-97.78, p=0.101). The production average of farmers having their ponds that are not overlapped with mangrove areas is 501.07 kg with a median 300 kg. While ponds that are overlapped with mangrove areas produce 429 kg with a median 700 kg, thus the discrepancy between them is only 1.6 kg.

**Fig 8.**
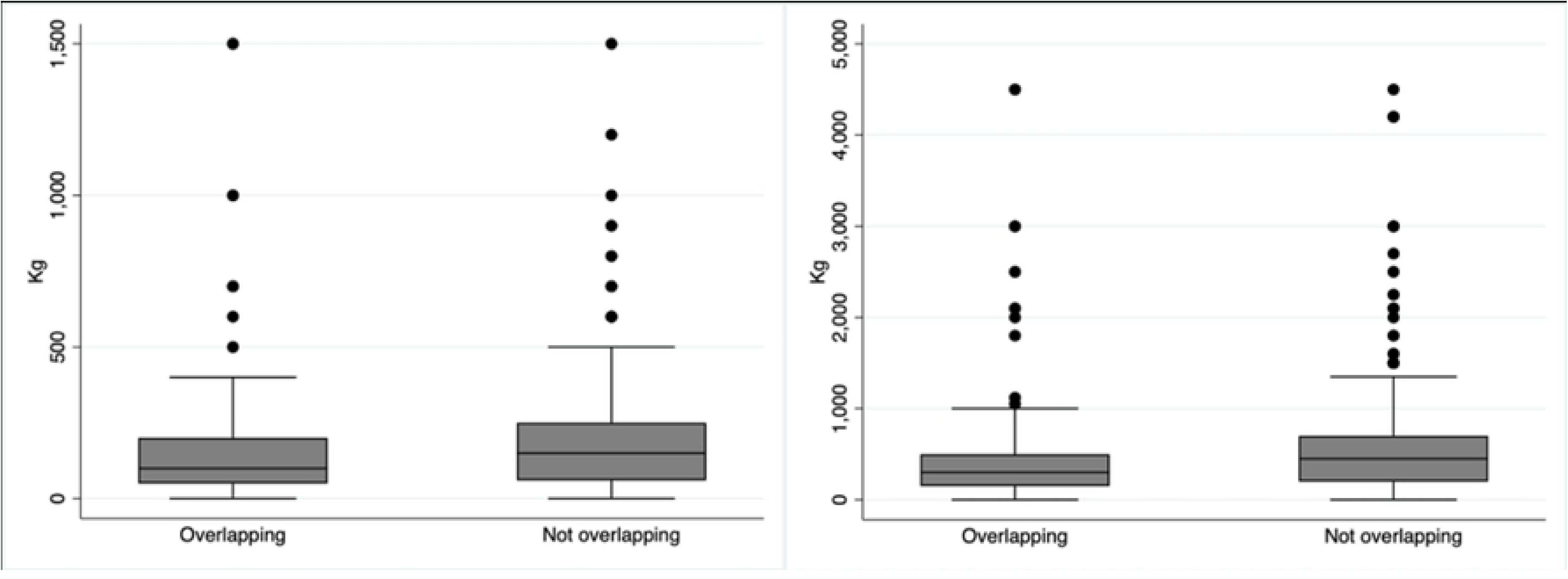
(Left) Total shrimp production in the last cycle based on external mangrove presence, N=619 (Right) Total shrimp production in 2022 based on external mangrove presence, N=619

The unequal t-test (see S7 Table) has shown that there is no significant difference between shrimp production from ponds overlapping and not with mangroves areas. Therefore, it is different with mangrove presence in the internal mangrove which depicts different shrimp production both in the last cycle and total in the whole year in 2022. As a further attempt, this study investigates the shrimp production’s difference between overlapped ponds with mangrove areas, which have varied mangroves densities, which were classified into low, medium, and high using Robust ANOVA.

Fig 9 shows a small difference production and revenue in the last cycle among density but the Robust ANOVA (see S8 Table) indicated that there is no significant of shrimp production in the last cycle (Brown-Forsythe’s F (2, 21.42) =1.26, p=0.301; Fisher’s F (2, 187) = 0.96, p=0.382; Welch’s F (2, 26.65) = 2.04, p=0.149). The discrepancy among overlapped mangrove density is only about 3 kg.

**Fig 9.**
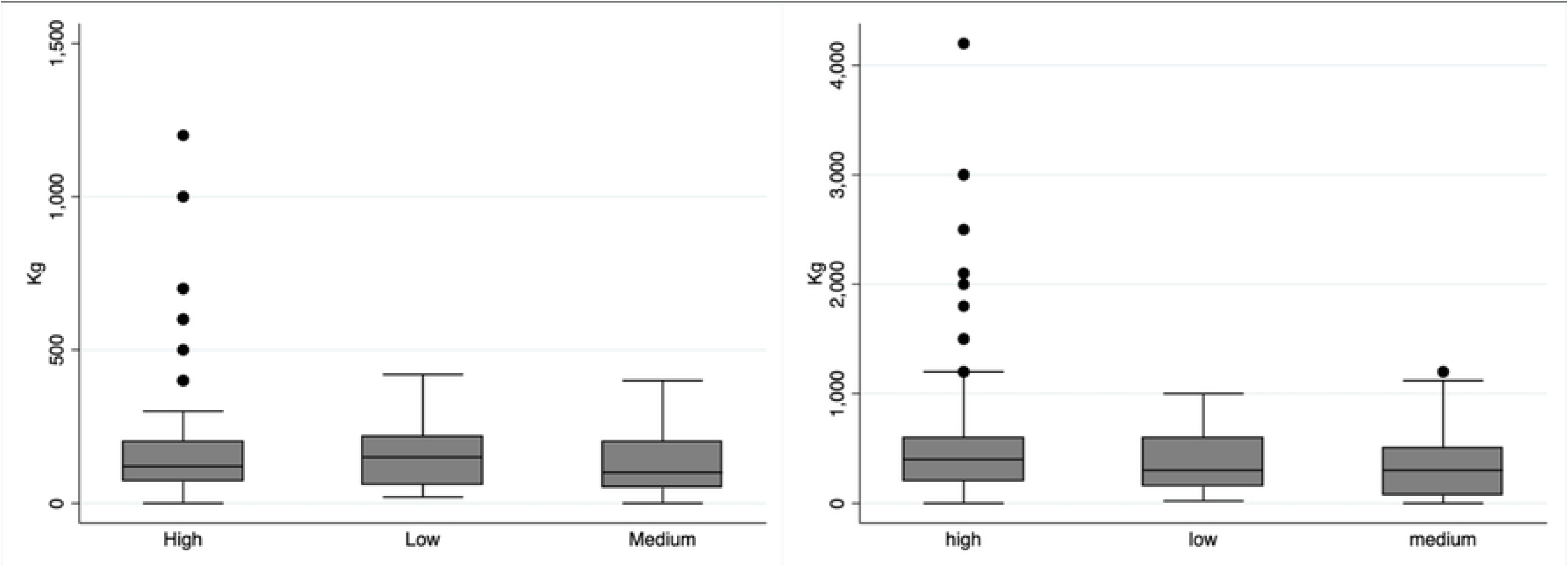
(Left) Total shrimp production in the last cycle based on external mangrove density, N=190 (Right) Total shrimp production in 2022 based on external mangrove density, N=190

### 3.3. Exploring Factors or Conditions Affecting Shrimp Production

The result of the first MANOVA results show that the simultaneous presence of internal and external mangroves does not significantly affect cyclical or yearly shrimp production. However, when assessed separately, internal mangrove presence has a significant negative effect on shrimp production for both cycles and yearly totals (Wilks’ lambda=0.96, p=0.000). Conversely, external mangroves alone do not significantly impact shrimp production (Wilks’ lambda=0.99, p=0.165).

As seen in S9 Table, only internal mangrove presence has a significant effect on the dependent variables (using Bonferroni Adjustment, tested at the 0.05 divided by 4 or 0,0125 level), total shrimp production in the last cycle and total shrimp production in 2022. The coefficient of internal mangrove presence is negative on cyclical and annual production. This indicates that mangrove presence has a negative relationship with the shrimp production. However, it should be noted that the R-squared of both total shrimp production in the last cycle and in 2022 is small, 0.023 and 0.041, implying that the internal mangrove presence explains only a small proportion of the variance of shrimp productions among ponds.

The result of the second MANOVA analysis, which examines mangrove position internally and the density of overlapped mangroves, indicates that these factors together affect both cyclical and yearly shrimp production (Wilks’ lambda = 0.91, F(6, 310) = 2.20, p-value = 0.042). Using the Bonferroni Adjustment, tested at the 0.05 divided by 6 or 0,008 level, there is a significant impact of the combination of internal mangrove position and external mangrove density on total shrimp production in the last cycle. It is a combination of having mangrove both inside and buffer in internal ponds and overlapping with high mangrove in external ponds (see S10 Table). They, together, have a significant positive impact on the last cycle of shrimp production. Other combinations have no significant effect due to factors such as limited sample size for each combination. It swayed the significance levels of particular category combinations. However, it’s advisable to investigate potential interaction effects and assess their implications comprehensively. Therefore, we can see that the ponds that with particular combination of characteristics tend to have a positive and a negative relationship with both total shrimp production in the last cycle and total shrimp production in 2022.

There are two conditions of the pond that have positive relationship with shrimp production, including 1) ponds having mangrove on both the inside and buffer, 2) ponds having overlapped mangrove with medium density outside of the pond boundaries. There are also the different conditions of the pond that have negative relationship with shrimp production, including the ponds having mangrove inside the ponds and having overlapped mangrove with medium density outside of the pond boundaries.

Specific effects of internal mangrove position also have a significant effect on the two dependent variables (Wilks’ lambda = 0.90, F(4, 310)= 3.94, p-value = 0.003). However, the impact of external mangrove density alone does not have a significant impact on shrimp production, both the cyclical production and the annual production. (Wilks’ lambda = 0.96, F(4, 310)= 1.31, p-value = 0.267). Ponds having mangrove both inside and buffering their ponds have a negative effect on shrimp production while ponds having mangrove inside the pond have positive effects on both cyclical and annual shrimp production. In addition, medium and high density also have negative effects on annual and cyclical production.

These two-way MANOVA results indicate that there are several conditions of mangrove affecting the shrimp production. The presence of mangrove inside the farmer’s ponds has a significant influence on shrimp production in both the last cycle and total production in 2022 (Table 2). The presence of mangrove in farmers ponds tends to decrease the shrimp production. On the other hand, the presence of mangrove in the external ponds does not influence the shrimp production. By observing the influence of internal mangrove position inside the farmers’ pond and the mangrove density around the farmer’s pond separately, the presence of mangrove inside and in the buffer negatively affects the shrimp production cyclical and annual. Similarly, the high-density mangrove around the ponds also negatively influenced the last cycle shrimp production.

**Table 2.**
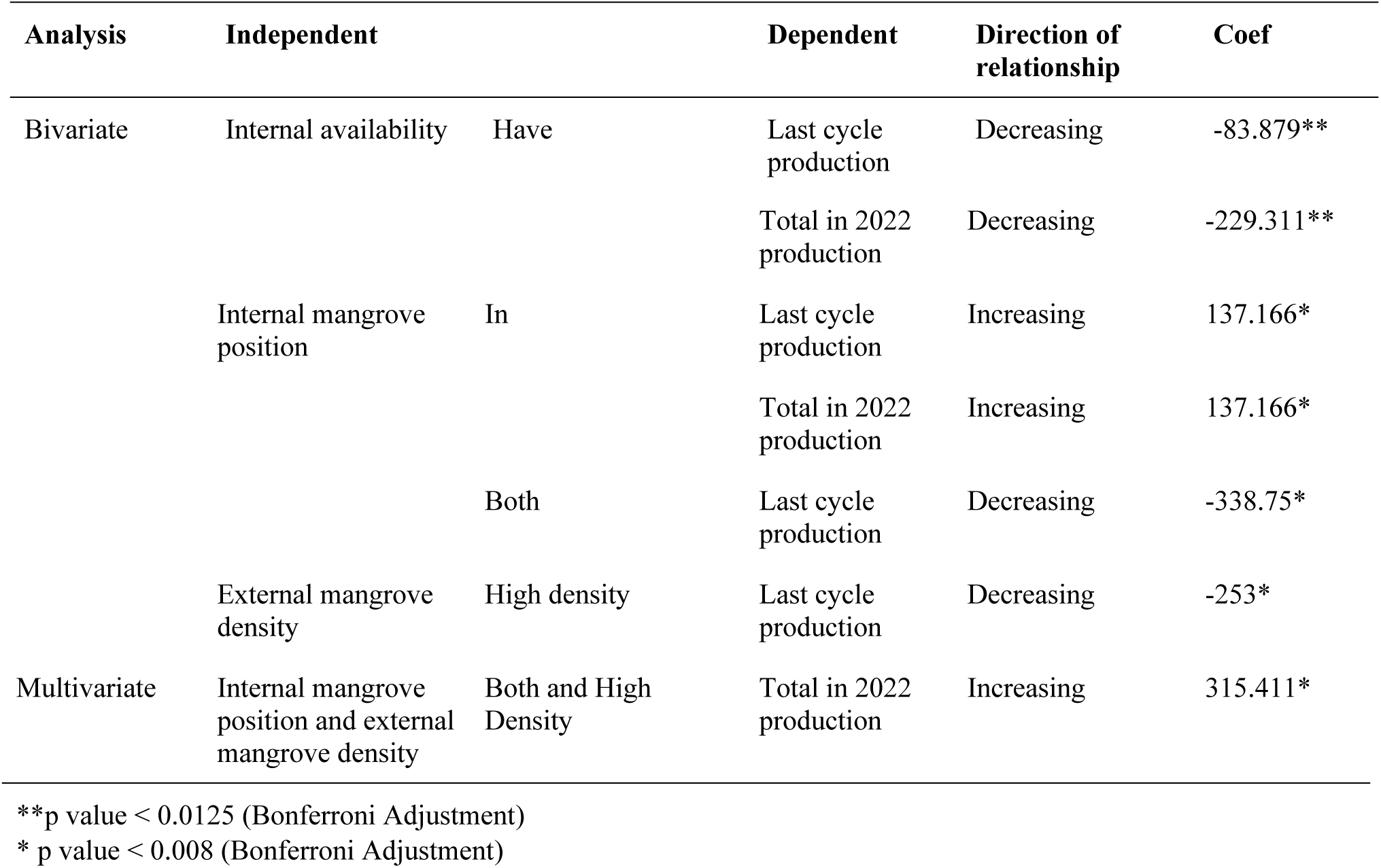
The relationship of mangrove on shrimp production (Two-Way MANOVA)

## Discussion

### Polarization of Perception Towards Mangrove and Shrimp Production

From both survey and interview, there is a polarization of perception and aspiration between shrimp farmers community in Berau in regards to mangrove in conjunction to shrimp aquaculture production. Majority of farmers see no correlation between mangrove and shrimp production. Some of the farmers perceive that mangrove have negative implications for the water quality, sources of diseases and predators, and overall reducing the shrimp production. Hence, for the farmers with negative perception, the pond should have been cleared from mangrove to improve the shrimp productivity. In contrast, some farmers perceive mangrove as integral to improve shrimp production by preventing land abrasion, providing food, nutrients, favorable microclimate, natural water filtering, and other services that improve shrimp production. The remaining farmers think that mangrove have both positive and negative relations to shrimp production. These farmers often mentioned that the inside of the pond is preferably clear from mangrove, meanwhile the wall of the pond can maintain the mangrove as a buffer that can maintain the structural integrity of the pond against extreme weather.

As indicated from the survey, the majority of farmers have mangrove within the pond boundaries, mainly because it is hard or costly for farmers to cut down and uproot the mangrove trees and there still some benefits of mangrove that farmers need for shrimp production. Some farmers with smaller plots of land also believe that they have limited production space, hence eliminating mangroves from the pond will increase the space needed for their shrimp production. As a result, these groups of farmers want to clear the pond from mangrove whenever they have opportunity or capital to do it because of a negative perception of low shrimp production owing to the existence of mangrove, impact of its litter, and its limited space.

Although the shrimp farmers recognize multiple factors affecting shrimp production, there are some farmers that still want to clear mangrove from their pond. From the interview, farmers have realized that there are multiple factors contributing to production of shrimp aquaculture such as shrimp seed quality, impact of extreme events (disaster, climate, and temperature changes), diseases, predatory attack, declining water quality within the overall water areas, having enough profit necessary to purchase high quality inputs for next cycle production, and good management practices (appointing experienced labor, regular litter and sedimentation clean up, and managing the use of fertilizers/chemicals). However, there is a stubborn belief that mangroves are counterproductive to shrimp aquaculture production.

Mangroves play a critical role in supporting sustainable shrimp aquaculture by providing ecosystem services that enhance water quality, reduce the impact of extreme weather events, and support biodiversity. Studies have shown that mangroves act as natural biofilters, improving water quality by trapping sediments and absorbing nutrients, which can mitigate the effects of declining water quality in shrimp ponds [29] Additionally, mangroves serve as a buffer against climate-induced disasters, such as storm surges and flooding, which are significant concerns for aquaculture [30]. Integrated mangrove aquaculture systems have been proposed as a sustainable alternative, demonstrating that mangroves can coexist with shrimp farming while maintaining ecological balance and improving long-term productivity [31]

### The Gap Between the Factual and Perceived Mangrove-Shrimp Production

In the bivariate analysis, there are three findings on the statistical analysis. First, this study finds that mangrove availability reduces both last cycle and annual shrimp production. Second, the statistical analysis also finds that mangrove positions inside the pond have a positive effect on both last cycle and annual shrimp production. For instance, mangroves positioned inside shrimp ponds can enhance water quality and provide natural habitats for shrimp, leading to improved production outcomes [32]. However, the presence of mangrove both inside and buffering the ponds have a negative effect on the shrimp production. Lastly, the high density of external mangrove forests has a negative effect on shrimp production. The negative effects of high-density external mangrove forests on shrimp production could be attributed to reduced water flow and increased organic matter accumulation, which may impact pond conditions [33]. Research also suggests that maintaining an optimal mangrove-to-pond ratio is crucial for balancing ecological benefits and aquaculture productivity [34].

In the multivariate analysis, there are two findings from the statistical analysis for shrimp production, internal mangrove position, and external mangrove density. After different statistical analysis, the author finds positive shrimp production in the year 2022 owing to the availability of mangrove both inside and buffer the pond and availability of high-density mangrove outside of the pond. The statistical analysis implies that mangrove conditions, such as availability, position, and density of mangrove in conjunction with a pond have different implications to shrimp production. The ability to comprehend the complex relation between mangrove and shrimp production is not accessible to the scientific community and local people. As an implication, there are circulating misconceptions toward mangrove and shrimp aquaculture relations. In addition, farmers also have to deal with extra burden to understand complicated factors that shape shrimp production, such as seed quality, extreme events, good management capacity, water quality, and other factors add more complexity for the shrimp farmers. Therefore, the mismatch between farmers’ perception of mangrove and its actual impact to shrimp production is anticipated due to lack of available information or evidence-based analysis shaping farmers’ negative perceptions.

### Possibility to Integrate Mangrove with Shrimp Aquaculture

From the quantitative analysis, mangrove and shrimp aquaculture is possible and supposedly lead to a productive relationship. However, there are two things that need to be considered for restoration implementers in regards to managing farmers’ perception and designing the restoration plan. Firstly, shrimp farmers’ negative perception of mangroves emerges from the live and personal experience that shrimp aquaculture is a livelihood activity which is sensitive to any environmental and even socio-economic changes [29, 31]. Therefore, any changes emerging from the successful or unsuccessful mangrove restoration efforts (such as integrating mangrove and shrimp aquaculture) can potentially be detrimental to the continuity of marginal shrimp farmers’ livelihood who are still productively farming [30]. Secondly, mangrove and shrimp aquaculture integration can be more strategic than planting mangrove seeds in and around the pond by analyzing thoroughly the type of mangrove species, the location of restoration, the intensity of the mangrove restoration, and overall distance and integration to the adjacent mangrove forest [35]. On the other hand, according to the farmers, integrating mangrove with shrimp aquaculture is not easy since the pond is a stable water without tidal changes exposure which might cause the death of mangrove plants. Anticipating the technical issue in mangrove restoration is also important for the sustainability of the project. With all these considerations in mind, the people involved in mangrove restoration activities need to consider multiple aspects including the continuity of livelihood in shrimp production and technicalities of mangrove restoration.

From the results, we develop scenarios for the implementation of IMSA. First, to maintain the shrimp production, the mangrove inside IMSA pond should be low in density if not absent. Second, if the IMSA pond is not productive or the production is low, there are options to adjust the mangrove density or fully restore ponds as intertidal mangrove patches. For pond full restoration, it is advisable to pursue restoration on the abandoned or non-productive shrimp farms as the low hanging fruits options. As a precaution, restoration efforts must also carefully consider mangrove density to prevent a production decrease in the neighboring ponds. Furthermore, fully incorporating the IMSA concept in Berau, especially regarding the tidal flows, might improve mangrove restoration projects. In addition, farmers’ aspiration for restoring mangrove-based coastal defense and mangrove buffer surrounding the pond for mitigating extreme events will be a great opportunity for restoration that is also the interest of the local community. To support the restoration effort, the implementation of IMSA should apply continuous monitoring to assess the quality and development of both the mangrove ecosystem and the aquaculture activities, ensuring sustainable management practices, identifying potential issues or challenges, and informing adaptive management strategies for long-term success [36].

### Management Implication

Our study demonstrates the theory of mangrove ecosystem services and disservices [5, 14] by showing that there is a polarization of perception among shrimp farmers in Berau regarding the role of mangroves in shrimp aquaculture production. Some farmers perceive them as beneficial, others as detrimental, while others acknowledge both positive and negative aspects of mangrove presence. While a majority of farmers fail to recognize any correlation between mangroves and shrimp production, some perceive mangroves as detrimental to water quality, disease control, and overall shrimp yields, advocating for their removal from ponds to enhance productivity. Conversely, another faction of farmers considers mangroves essential for improving shrimp production by preventing land abrasion, providing essential nutrients, maintaining a favorable microclimate, and offering natural water filtering. Meanwhile, others acknowledge both positive and negative aspects of mangrove presence, suggesting clearance from inside ponds while maintaining them along pond walls as protective buffers. Despite recognizing various factors influencing shrimp production, including seed quality and management practices, some farmers continue to believe that mangroves hinder shrimp aquaculture, highlighting the complex and contentious nature of perceptions within the farming community.

This study sheds light on the relationship between mangroves and shrimp production, offering insights into the discrepancies between farmers’ perceptions and empirical evidence. It advances the research of perception on mangrove and shrimp [19] and empirical study of mangrove relationship with shrimp production [20]. Despite farmers’ negative perceptions of mangroves, our findings suggest that mangroves can have a positive effect on shrimp production when considering their position and density. While farmers’ negative perceptions of mangroves may stem from their observations of shrimp production in ponds with and without mangroves, the analysis reveals that mangroves within ponds tend to reduce shrimp production, while their presence in buffering areas can also have negative effects. However, simultaneous consideration of mangrove position and density indicates that having mangroves both inside and buffering the ponds, along with higher mangrove density outside the ponds, correlates positively with shrimp production. This contradicts the common perception among the community that clearing mangroves within and around ponds is beneficial, as the statistical findings demonstrate that the presence of mangroves within ponds, surrounding them, and in high-density forest nearby actually correlates positively with shrimp production. Thus, addressing the perception is crucial to support community livelihood and enhance climate mitigation via mangrove conservation and restoration initiatives.

While there is potential for a productive relationship between mangrove and shrimp aquaculture, negative perceptions often obstruct their integration. The community holds the belief that mangroves detrimentally affect shrimp production, despite circumstances where mangroves might enhance it. To address this, restoration implementers must consider anticipating livelihood disruptions [15,18] and strategically planning the integration as well managing technical challenges [21]. These considerations are essential for overcoming obstacles that come from the community and ensuring the success of mangrove restoration efforts are in line with socio-economic well-being and are in line with socio-economic well being. It highlights that IMSA is not a universal solution where its effectiveness varies depending on various conditions and needs thorough and routine monitoring and evaluation.

Our recommendation stems from the expansion of the argument of Lovelock et al. [12] on mangrove restoration. Our study proposes scenarios of the implementation of IMSA, including adjusting mangrove density and restoration of non-productive or neglected ponds by considering the sustainability of pond production in the entire area and ecosystem. Further to this, refer to the priority location for mangrove restoration in Lovelock et al’s research [12], implement appropriate tidal water exchange in the shrimp aquaculture system could be a considerable measure to achieve IMSA’s goals in Berau, but the impact on shrimp production for the farmers need further clarification.

Finally, it is important to acknowledge that there are numerous other factors influencing production that were not analyzed in this study, such as the type of native or non-native species, the distribution and type of mangrove species, and the presence of other species like milkfish and crabs. These factors present opportunities for further research (for example, productivity function in shrimp aquaculture research that combine multiple factors for shrimp productivity) [37]. Additionally, studying the impact of IMSA on livelihood disruption could also be a valuable avenue for future research.

## Acknowledgments

This research was supported by multiple funding including the Darwin Initiative (project title: Nature Climate Solution to Protect Mangrove Biodiversity and Improve Livelihood--project reference 29-029) and The Nature Conservancy-NCS Prototyping Network Grant. The fundings was instrumental in supporting field data collection, analysis, and manuscript preparation. We would also like to express our deepest gratitude to the local communities of Pegat Batumbuk, Suaran, Tabalar Muara, Sukan Tengah, Pilanjau, and Gurimbang for their invaluable support during fieldwork. Their willingness to share their knowledge, experiences, and hospitality greatly enriched this research. We are particularly thankful to the community leaders and enumerators —Indra, Marina, Dewi, Natan, Hasnur, Milov, Walhiswandi, Udi, and Windi —for their dedication in conducting interviews, facilitating discussions, preparing logistics, and ensuring the success of this study. This research would not have been possible without their cooperation and insights.

## Author contributions

AWA: Conceptualization, Data curation, Resources, Project administration, Writing – review & editing; YO: Conceptualization, Data curation, Formal analysis, Methodology, Visualization, Project administration, Writing – original draft; DAN: Data curation, Formal analysis, Visualization, Writing – original draft; NDA: Data curation, Formal analysis, Methodology, Writing – original draft; Muhammad Ilman: Supervision, Writing – review & editing. All authors: Writing – review & editing.

## Supporting Information

**S1 Table.** Distribution of survey data based on village

**S2 Table.** Pond-Level Characteristics and Associated Respondent Demographics (N=619)

**S3 Table.** In-depth interview coding system

**S4 Table.** Bartlett’s equal of variances test

**S5 Table.** Unequal t-test for shrimp production difference based on mangrove availability

**S6 Table.** Robust ANOVA of shrimp production based on mangrove presence in internal ponds

**S7 Table.** Unequal t- test shrimp production based on overlapped and not overlapped

**S8 Table.** Robust ANOVA of shrimp production and revenue, based on overlapped mangrove density

**S9 Table.** Effect of internal and external mangrove presence on shrimp production (N=619)

**S10 Table.** Effect of mangrove presence on shrimp production (N=164)

## Notes

### Competing Interest Statement

The authors have declared no competing interest.

